# A vimentin-targeting oral compound with host-directed antiviral and anti-inflammatory actions addresses multiple features of COVID-19 and related diseases

**DOI:** 10.1101/2021.08.26.457884

**Authors:** Zhizhen Li, Jianping Wu, Ji Zhou, Baoshi Yuan, Jiqiao Chen, Wanchen Wu, Lian Mo, Zhipeng Qu, Fei Zhou, Yingying Dong, Kai Huang, Zhiwei Liu, Tao Wang, Deebie Symmes, Jingliang Gu, Eiketsu Sho, Jingping Zhang, Ruihuan Chen, Ying Xu

## Abstract

Damage in COVID-19 results from both the SARS-CoV-2 virus and its triggered overreactive host immune responses. Therapeutic agents that focus solely on reducing viral load or hyperinflammation fail to provide satisfying outcomes in all cases. Although viral and cellular factors have been extensively profiled to identify potential anti-COVID targets, new drugs with significant efficacy remain to be developed. Here, we report the potent preclinical efficacy of ALD-R491, a vimentin-targeting small molecule compound, in treating COVID-19 through its host-directed antiviral and anti-inflammatory actions. We found that by altering the physical properties of vimentin filaments, ALD-491 affected general cellular processes as well as specific cellular functions relevant to SARS-CoV-2 infection. Specifically, ALD-R491 reduced endocytosis, endosomal trafficking, and exosomal release, thus impeding the entry and egress of the virus; increased the microcidal capacity of macrophages, thus facilitating the pathogen clearance; and enhanced the activity of regulatory T cells, therefore suppressing the overreactive immune responses. In cultured cells, ALD-R491 potently inhibited the SARS-CoV-2 spike protein and human ACE2-mediated pseudoviral infection. In aged mice with ongoing, productive SARS-CoV-2 infection, ALD-R491 reduced disease symptoms as well as lung damage. In rats, ALD-R491 also reduced bleomycin-induced lung injury and fibrosis. Our results indicate a unique mechanism and significant therapeutic potential for ALD-R491 against COVID-19. We anticipate that ALD-R491, an oral, fast-acting, and non-toxic agent targeting the cellular protein with multipart actions, will be convenient, safe, and broadly effective, regardless of viral mutations, for patients with early- or late-stage disease, post-COVID complications and other related diseases.

**IMPORTANCE:** With the Delta variant currently fueling a resurgence of new infections in the fully-vaccinated population, developing an effective therapeutic drug is especially critical and urgent in fighting COVID-19. In contrast to the many efforts to repurpose existing drugs or address only one aspect of COVID-19, we are developing a novel agent with first-in-class mechanism-of-actions that address both the viral infection and the overactive immune system in the pathogenesis of the disease. Unlike virus-directed therapeutics that may lose efficacy due to viral mutations and immunosuppressants that require ideal timing to be effective, this agent, with its unique host-directed antiviral and anti-inflammatory actions, can work against all variants of the virus, be effective during all stages of the disease, and even resolve post-disease damage and complications. A further development of the compound will provide an important tool in the fight against COVID-19, its complications, as well as future outbreaks of new viruses.

## INTRODUCTION

Vaccines against SARS-CoV-2 have been rapidly developed and rolled out to contain the COVID-19 pandemic, however the infection-blocking immunity induced by vaccination has been shown to wane quickly, even though disease-reducing immunity may be long-lived (1). Vaccination primarily generates IgG antibodies that circulate in the blood, not IgA antibodies in the mucosa of the respiratory tract which are a critical component of the mucosal immunity that prevents SARS-CoV-2 from initiating its infection (2, 3). The spike mRNA-based vaccination induces strong, persistent antibody responses but accompanied by a limited or absent T-cell response (4, 5). In addition, the virus has now demonstrated an ability to rapidly mutate during its transmission and to generate mutants that not only provide enhanced infectivity, but also an acquired ability to escape from therapeutic antibodies and those induced by infection or vaccination. Several Variants of Concern (VOCs) have emerged within a short period of time, especially the Delta variant (6, 7) that is currently fueling a resurgence of new infection in fully vaccinated population. This virus appears to present a persistent problem despite vaccines, and therefore effective therapeutics are critically needed.

To date, most therapeutic interventions have been focused either on stopping viral infection or reducing virus-induced hyper-inflammation. Antiviral approaches can be virus-directed or host-directed. Virus-directed approaches include blocking viral entry, preventing maturation of viral proteins, and disrupting the viral RNA-synthesis machinery. All face challenges from the rapid viral mutations that could make them ineffective over time. In addition, these types of agents must be given early during an infection to be effective, as viral load has been shown to peak on or before symptom onset (8) with very few infectious viruses present in the upper respiratory tract 10 days after the onset of symptoms, even without treatment (9). A delayed or longer anti-viral treatment in the course of COVID-19 has been shown to lead a worse outcome (10), because it is the overactivated host immune system, not the virus itself, that is driving the progression and the development of serious complications. However, immunomodulators have shown minimal effects and corticosteroids, moderate effects only if given at the right timing, otherwise worsening the disease. Host-directed antiviral approaches have attracted attention because targeting host proteins offers broad-spectrum anti-viral actions that are hard for virus to evade through mutation. The host proteins that interact with SARS-CoV-2 have been extensively profiled and a large quantity of existing compounds have been screened for potential antiviral activities (11–15). So far, however, no repurposed antiviral drugs addressing these host factors have shown significant efficacy in clinical trials(16).

An ideal anti-COVID agent would be able to both eliminate the virus and restore balance to the immune system. Several existing drugs, such as hydroxychloroquine, colchicine, and ivermectin that exhibit some degree of synergistic antiviral and anti-inflammatory actions, have been repurposed to treat COVID-19, however, the results have been controversial or inconclusive. Whether applied in early disease to prevent progression, or in late stage to reduce morbidity and mortality, therapeutics with new mechanisms are urgently needed to tackle the multifaceted and inter-related nature of pathogenesis in SARS-CoV2 infection.

Vimentin, an evolutionarily very conserved intermediate filament protein with diverse functions in health and disease (17), has been proposed as a host-directed therapeutic target for COVID-19 (18, 19). Vimentin facilitates viral infection by serving as a co-receptor and a component of the cellular transportation machinery for viral entry, trafficking and egress (20). Vimentin is also specifically involved in inflammatory responses, where it forms a superstructure inside regulatory T cells that mechanically regulate the switch of the cells between active and inactive status (21), and provides a scaffold structure that enables the NLRP3 inflammasome to assemble, activate, and contribute to inflammation and lung injury (22). Targeting vimentin is unlikely to generate mechanism-based toxicities because the protein has been widely recognized to be non-essential from embryonic development to adulthood after the first work in early 1990s demonstrated that mice lacking vimentin developed and reproduced without an obvious phenotype (23).

We synthesized a series of small molecule compounds that can bind to vimentin (24). The lead compound ALD-R491 has shown many favorable safety features: it is fast-acting, fully reversable, has no cytotoxicity, and causes no change in the protein levels of vimentin and other proteins of major signaling pathways (25). In this study, we explored the effects of ALD-R491 on vimentin structure and function, general cellular processes and specific cellular functions relevant to SARS-CoV-2 infection, and evaluated the compound *in vitro* and *in vivo* for its antiviral and anti-inflammatory effects. The constellation of effects we observed strongly supports this oral small molecule compound as a promising host-directed therapeutic agent with novel mechanisms to address multiple pathological processes of COVID-19.

## RESULTS

### ALD-R491 reduces the dynamism of vimentin filaments and inhibits endocytosis, endosomal trafficking and exosomal release

To understand the impact of the compound-binding on the target protein, we first examined the distribution of vimentin intermediate filaments (IF) in live cells. Confocal immunofluorescence imaging showed a dynamic reorganization of intracellular vimentin network responding to ALD-R491 treatment. In the control cells, the vimentin IF formed bundle-like structures parallel to the longitudinal axis of the cell, with a richer presence in the cell peripheral region and at the far end of cell than in the central region. In ALD-R491-treated cells, the vimentin IF retracted from the periphery with no structural changes visible at low doses but with an intricate, honeycomb-like reticulated structure formed in the perinuclear region at high doses (**Fig. 1A**). This alteration was specific to vimentin, because the proteins of the same type-III IF family, GFAP and desmin, showed no structural changes to ALD-R491 treatment. (**Fig. S1A**).

**Fig. 1.**
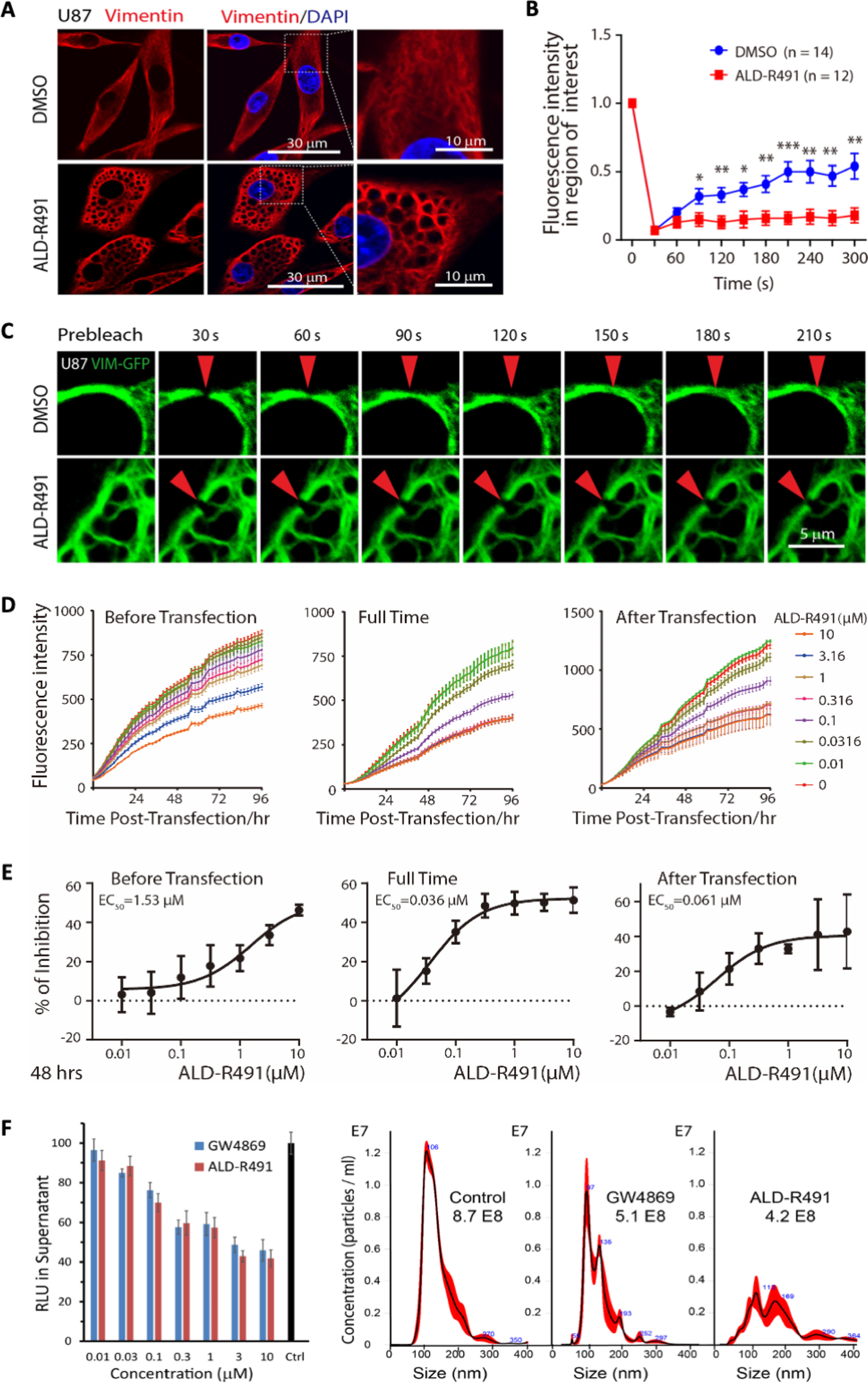
Effects of ALD-R491 on vimentin filaments and sub-cellular processes. changes in intracellular organization and physical property of vimentin filament and reductions in endocytosis, endosomal trafficking and exosomal release. **A.** ALD-R491 induced the reorganization of vimentin intermediate filament network. **B-C.** ALD-R491 compromised the mobility of vimentin filaments. **B.** Quantitative measurement; **C.** Image presentation. Two-tailed T Test. * p<0.05, ** p<0.01, *** p<0.001. **D-E.** Dose-responsive inhibition on endocytosis and endosomal trafficking. **D.** Fluorescence intensity over time; **E.** Dose-responsive reduction in GFP levels. **F.** Exosome release blockade. Left bar graph: the luciferase activity in the culture supernatant was quantified; Right: the purified exosomes were quantified by nanoparticle tracking analysis.

We next sought to determine how the altered organization of vimentin IF would relate to a change in the physical properties of ALD-R491-bound vimentin filament. We utilized the fluorescence recovery after photo-bleaching (FRAP) technique to quantify the two-dimensional lateral diffusion of vimentin-GFP in the cells treated with the compound or the solvent control. We found that the time required for fluorescence recovery in the photo-bleached region was significantly longer in the ALD-R491 treated cells (**Fig. 1B, 1C**), indicating that the ALD-R491-bound vimentin protein had become less mobile or had a compromised dynamism.

Vimentin is involved in the trafficking of membrane and intracellular vesicles, so we hypothesized that a decrease in mobility or dynamism would impact subcellular processes related to material transportation. To test this, we used a liposome-mediated plasmid transfection method. To separate the effect on endocytosis from that on endosomal trafficking, we treated the cells with ALD-R491 during three different time periods (**Fig. 1D-E**). When present during the entire transfection process (full time) that impacts both endocytosis and trafficking, ALD-R491 reduced the levels of GFP reporter up to 50%, with an EC50 of 0.036 μM and EC90 of 0.19 μM. At 0.3 μM ALD-R491 reached its maximum effect of 50% inhibition (treated full time), and the inhibition rates of ALD-R491 on endocytosis (treated before transfection) and endosomal trafficking (treated after transfection) were 18% and 35% respectively (**Fig. 1E**). These results indicate that endocytosis and endosomal trafficking were very sensitive to ALD-R491 however neither of cellular processes was completely blocked by the compound, with endosomal trafficking impacted more than endocytosis.

Exosomes originate from the endocytic pathway and are released via the fusion of a multivesicular body (MVB) with cell membrane (26). Using a reporter system as well as a classic exosome purification method, we found that ALD-R491 inhibited exosome release from Huh7 cells in a dose dependent manner (**Fig. 1F**). Like the effects observed on endocytosis and endosomal trafficking, the maximal rates of inhibition on exosome release were plateaued at around 50% of non-drug levels. These data indicate that by altering the mechanical properties (“dynamism”) of vimentin filaments, ALD-R491 reduced multiple cellular processes that are related to viral entry, trafficking and egress.

### ALD-R491 blocks SARS-CoV2 spike protein-ACE2-mediated pseudoviral infection

Targeting the initial steps of infection, such as viral entry and de-coating, is an attractive antiviral approach, particularly for a host-directed antiviral mechanism with the objective of blocking virus-host cell interactions. SARS-CoV-2 gains access into cells through endocytosis after the engagement of its spike protein with cell membrane proteins. Angiotensin-converting enzyme 2 (ACE2) is the receptor for the spike protein and is responsible for the initiation of infection, although other host factors such as TMPRSS-2 and neuropilin-1 may facilitate the viral entry process. We tested the antiviral activity of ALD-R491 by using a lentivirus pseudotyped with the human SARS-CoV-2 spike protein to infect HEK293 cells overexpressing human ACE2 protein.

Because the pseudovirus expresses both GFP and luciferase, the infection efficiency can be continuously tracked by live imaging of fluorescence and quantified at the endpoint by measuring luciferase activity. Exposure of the cells to ALD-R491 resulted in a dramatic reduction in the pseudoviral infection at all dose levels during the whole infection process (**Fig. 2A-B, Fig. S2A-B**), and generated no cytotoxicity even at the highest concentration tested (CC50 >10 µM) (**Fig. 2C**). We conclude therefore that this antiviral effect was largely due to a blockade of viral entry rather than subsequent steps after endocytosis because no antiviral activity was observed when the compound was given 2 h after the viral infection. Regardless of the level of viral load present, ALD-R491 strongly and dose-dependently inhibited pseudoviral infection, with IC50s of 13.5, 34.7 and 64.9 nM for the infections at MOI of 0.5, 5 and 50, respectively (**Fig. 2D-E**).

**Fig 2.**
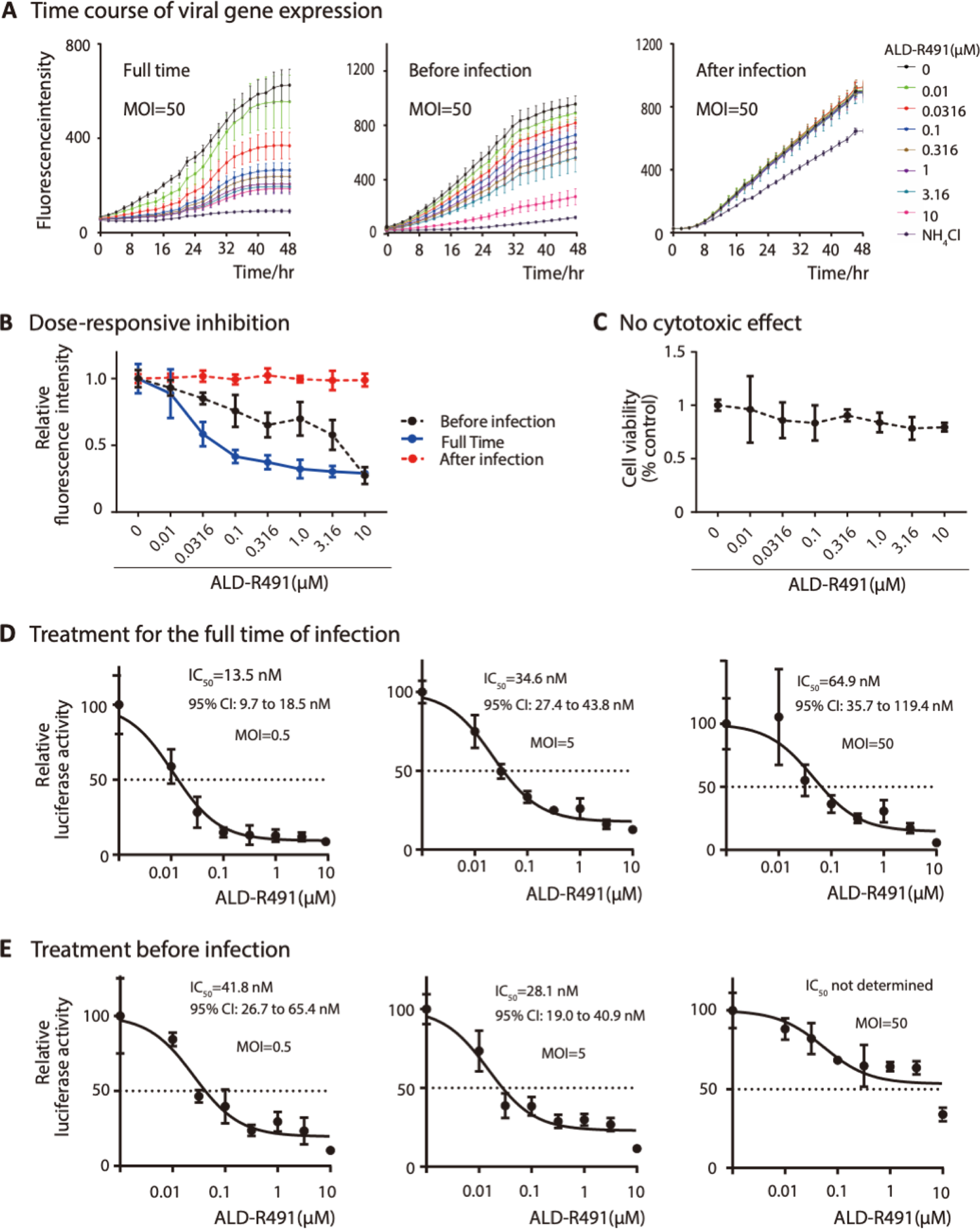
ALD-R491 blocked Spike protein-ACE2 mediated viral entry *in vitro.* **A.** Fluorescence intensity of the GFP reporter gene expression. The HEK cells were infected by the GFP-expressing lentiviral-based pseudovirus and measured for fluorescence intensity every 2 h over the course of the experiment. The cells were treated with ALD-R491 from 2 h before the infection till the end of the experiment (Left, Full time), for only 2 h before the infection with the removal of the compound upon the infection (Center, Before infection) and from 2 h after the infection till the end of the experiment (Right, After infection). **B.** Dose-dependent GFP levels at 48 h after infection presented. **C.** No cytotoxicity associated with ALD-R491**. D-E.** At the end of the experiment, luciferase activities were measured from the cell lysates. The figures show the inhibition rates of ALD-R491 at different concentrations on the infection by the pseudovirus at the multiple of infection (MOI) of 0.5, 5 and 50 respectively. Exposure of cells to ALD-R491: **D.** Full time; **E.** Before infection. IC_50_ and its 95% CI shown in the figures.

Clearly, ALD-R491 was highly selective (> 154 at MOI of 50) and potent at blocking the spike protein-ACE2-mediated viral entry, the first step of SARS-CoV2 infection. Unlike virus-directed approaches aimed at blocking the binding of the spike protein with ACE2 receptor, which include anti-spike protein antibodies that may become futile as the epitope of spike protein continues to mutate, our host-directed approach blocks the cellular process that is used by the virus. As long as endocytosis remains the route for their cellular entry, the compound would be equally effective for all variants.

### ALD-R491 enhances macrophage’s pathogen-killing efficiency

Monocytes and macrophages are the most important innate immune cells in the host defense against infections from pathogens, including viruses, fungi, and bacteria. At the early stages of an infection, the innate immune system, in particular macrophage M1, is as the first line of defense that prevents pathogens that have entered the tissues from further dissemination. In addition to triggering an immune response and producing proinflammatory cytokines, macrophages effectively phagocytose these pathogens, directly destroying them inside the phagosome or lysosome. SARS-CoV2 can interrupt phagolysosome function and even hijack the phagolysosome to egress from infected cells (27). The SARS-CoV-2 virus has been found alive inside monocytes and macrophages (28, 29), suggesting an impaired function of these cells for pathogen clearance, and a problematic reservoir site that supports lingering infection. The key actor for macrophage microbicidal activity is NADPH oxidase in the phagosomes which produces reactive oxygen species (ROS). NADPH oxidase is a membrane-associated enzyme consisting of multiple subunits. Vimentin is known to bind with p47Phox, a subunit of NADPH oxidase. The association with vimentin makes the p47phox subunit separated from the enzyme complex of NADPH oxidase, restraining the enzyme activity thereby limiting ROS production and consequently reducing the pathogen-killing efficiency of cells (30). In the mouse peritoneal macrophages, we found that p47Phox was abundantly present in the cell periphery next to the cell membrane and colocalized with vimentin. The vimentin-binding action of ALD-R491 caused a dissociation of vimentin from p47Phox thereby increased the presence of p47Phox in the cytosol (**Fig. 3A-B, Fig. S3**), increased the production of ROS (**Fig. 3C**) and reduced the number of Salmonella survived in the macrophages by about 30% (**Fig. 3D**). These results suggest a potential of ALD-R491 to increase the pathogen-killing capacity of macrophages. As viral evasion of host immune surveillance is believed to play a major role in disease severity and persistence, the enhanced microcidal function of macrophage by ALD-R491 could reduce the ability for SARS-CoV2 to survive once inside the macrophage, thereby decreasing the chance of viral persistence and preventing the escalation of inflammation and development of long-haul symptoms.

**Fig 3.**
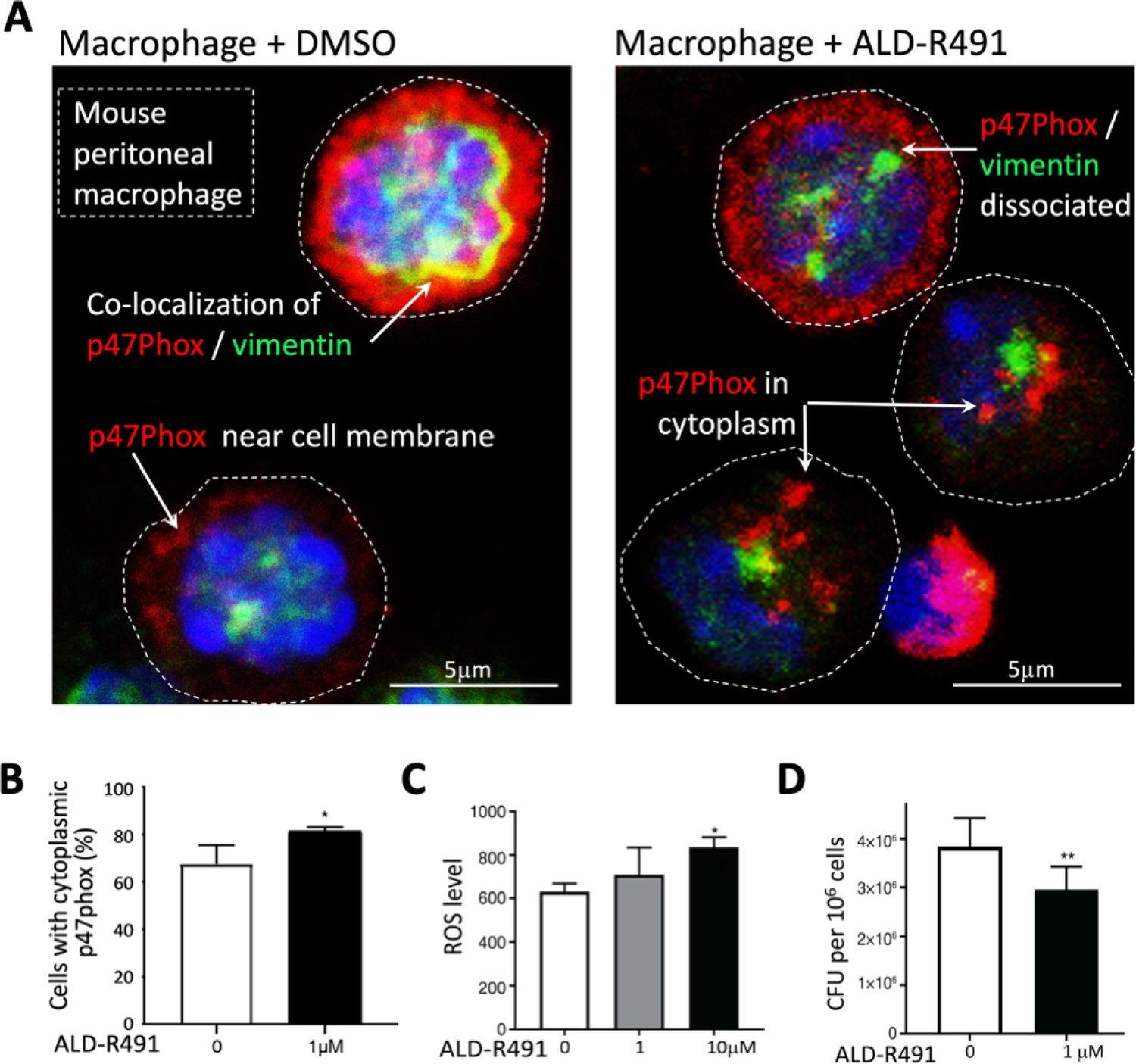
Effects of ALD-R491 on macrophages *in vitro*. **A-B.** ALD-R491 dissociated vimentin from p47Phox of NADPH oxidase in mouse peritoneal macrophages: **A.** Representative Images of macrophages with or without ALD-R491 treatment; **B.** Quantitation of p47phox dissociated from vimentin from 3 independent experiments. **C**. ALD-R491 increased cellular production of reactive oxygen species from Raw246.7 cells. from three independent experiments. **D.** ALD-R491 reduced the numbers of live Salmonella inside Raw246.7 cells, from six independent experiments. Two tailed T Test. * p<0.05, ** p<0.01.

### ALD-R491 directly activates regulatory T cells

In cases where the innate immunity fails to clear the pathogen, the adaptive immunity is then mobilized against the infection. Although this will help control infection, the overactive immune response is the most important contributor to morbidity and mortality in severe COVID-19. Among all immune cells, regulatory T cells (Tregs) are the key enforcers of immune homeostasis, marshalling many other immune cells in concert to keeping inflammation in check. They prevent host tissues from damage from the pathogen-triggered excessive antimicrobial immune responses and promote healing to repair epithelial damage, and other tissue damage throughout the body. In severe COVID-19 patients, the levels of peripheral Tregs were found to be remarkably reduced (31–34), which has led to a call for Treg cell-based therapy (35) and clinical trial (NCT04468971) of infusions of Treg cells. Vimentin has been previously identified as forming the ‘mechanical’ restraint structure inside regulatory T cells that regulate their switch between active and inactive status. Specifically, inside Tregs, the asymmetric partitioning of vimentin forms the distal pole complex (DPC), keeping the immunosuppressive molecules away from the immunological synapse that gets formed between Tregs and antigen-presenting cells, thereby restraining the function of Treg cells (Fig. S3B) (21). We hypothesized that ALD-R491 would interrupt the DPC’s structural integrity and directly activate Tregs to increase their dampening and balancing actions, thereby addressing and potentially solving the cytokine storm of COVID. In cultured mouse Treg cells treated with ALD-R491 for a short period of time (2 h), concentrations as low as 0.01 mM resulted in a significantly decreased proportion of inactive Treg cells that had intact DPC (**Fig. 4A-B**). When disassembled, DPC would release the immunosuppressive molecules, activating Treg cells. Indeed, Treg suppression assay in vitro showed an increased Treg activity in the presence of ALD-R491 at a low concentration of 0.1 µM (**Fig. 4C, Fig. S4**). These results indicated that the highly ordered DPC structure was very sensitive to ALD-R491 and Treg cells required very low concentrations of the compound to be activated.

**Fig 4.**
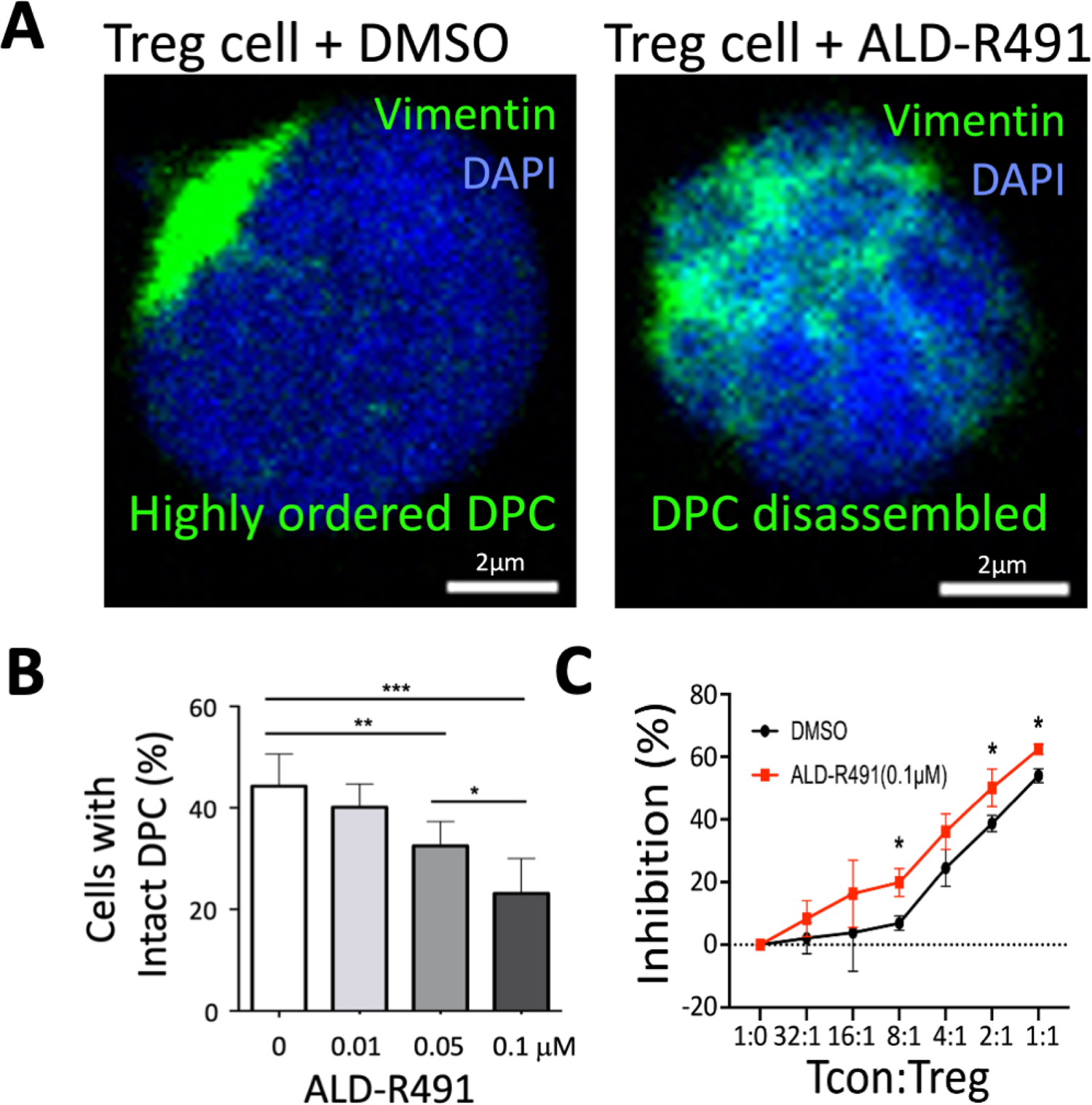
Effects of ALD-R491 on regulatory T cells *in vitro*. **A.** ALD-R491 disassembled the distal pole complex (DPC) of mouse regulatory T cells. Representative images of Treg cells with or without ALD-R491 treatment; **B.** Quantitative analysis of Treg cells with disassembled DPC, from three independent experiments. **C.** Quantitative measurement of Treg cell activity, from three independent experiments. Two tailed T Test. * p<0.05, ** p<0.01, *** p<0.001.

### Orally administered ALD-R491 shows therapeutic efficacy against SARS-CoV2 infection in aged mice

Both macrophages and Tregs are highly heterogeneous and plastic, and their functions *in vivo* are largely context dependent. We focused on whether their enhanced functions by ALD-R491 observed *in vitro* would translate into therapeutic benefit against COVID-19 *in vivo*. To evaluate whether the oral compound would have exposure in the lung tissue at levels sufficient to be effective, we first measured the distribution of ALD-R491 in tissues after oral administration. In rats gavaged with ALD-R491 at 30 mg/kg, the compound was rapidly distributed in various tissues, with the highest exposure in the gastrointestinal tissues as expected, enriched exposures in other tissues including the lung and lymph nodes, and minimal exposure in the blood (**Fig. 5A**). The compound concentrations in the lung tissue 6 h after oral dosing maintained at levels as high as 1μM, which was a concentration more than sufficient for efficacy based on its effective concentrations *in vitro* (< 0.1μM in multiple assays). These data provided the rationale for dose selection and supported a daily dosing treatment regimen.

**Fig 5.**
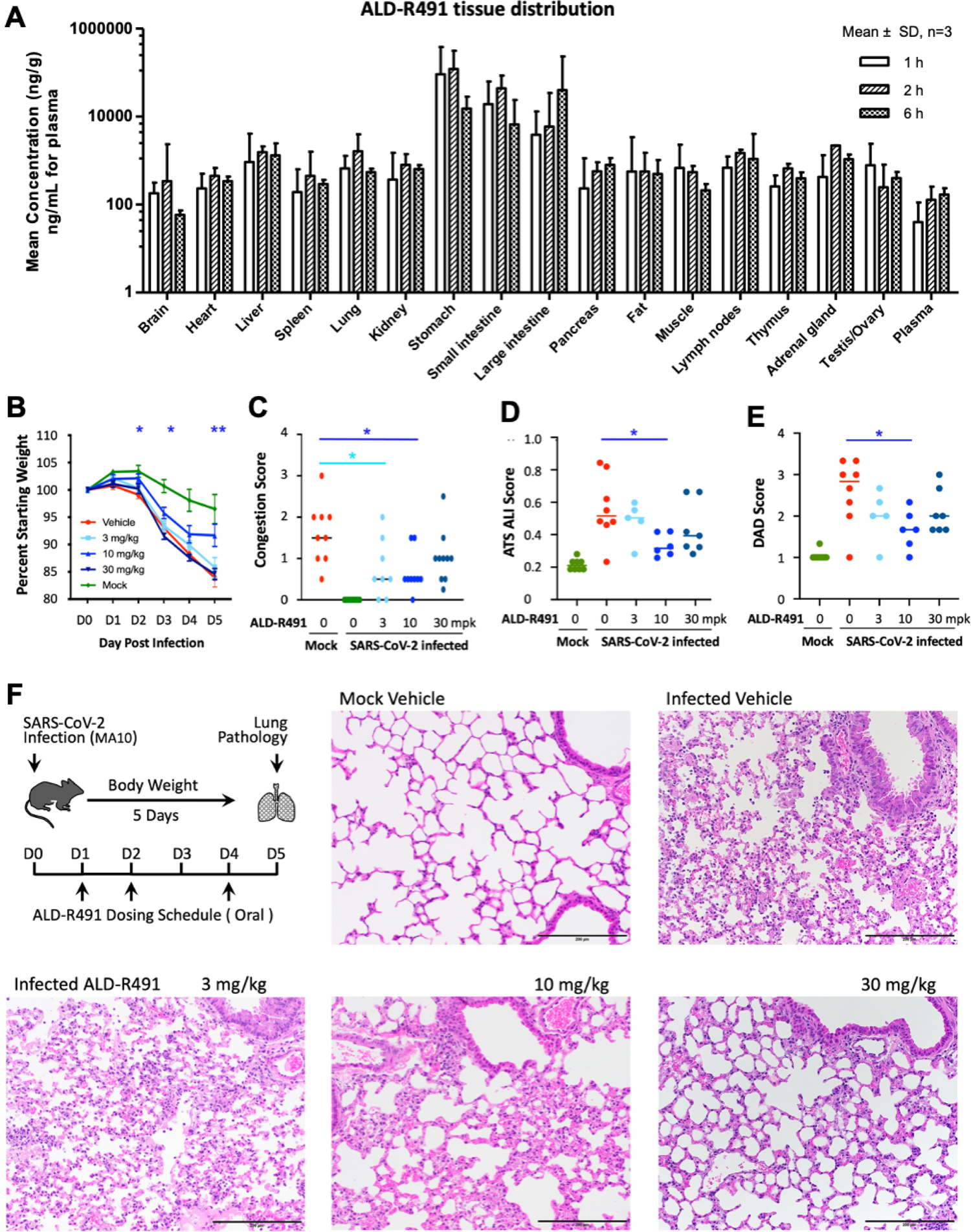
ALD-R491 tissue exposure in rats and therapeutic efficacy against SARS-CoV-2 infection in aged mice. **A.** ALD-R491 concentrations were measured in the indicated tissues of female rats 1 hr, 2 hrs and 6 hrs after oral administration of the compound. **B-F.** Therapeutic efficacy of ALD-R491 in aged mice Balb/c mice infected with SARS-CoV-2 M10. Body weights were measured daily over the course of study. Macroscopic and microscopic presentations of the lung at necropsy and in histopathology examination were blindly evaluated by a board-certified pathologist. The scores were compared between the Infected Vehicle group vs. individual treatment groups. Two tailed T Test, * p<0.05, ** p<0.01. **B.** Body weight change. **C.** Lung congestion scores on Day 5. **D.** ATS Acute Lung Injuries (ALI) scored by the method of Matute-Bello (2011). **E.** The diffuse alveolar damage (DAD) scored by the method of Schmidt (2018). **F**. Schematic presentation of the experimental protocol and representative images selected from mice with the ALI and DAD scores close to the scores of their group averages.

Next, we tested the compound in a mouse-adapted SARS-CoV-2 MA model, in which the mutated SARS-CoV-2 virus can replicate in the upper and lower airways of both young adult and aged BALB/c mice. The model generates more severe disease in aged mice, reproducing the age-related increase in disease severity observed in humans (36). In this study, we utilized aged mice (11 to 12 months) to best mimic the most severe viral infection in the most vulnerable population in humans. The compound was orally administered prophylactically and therapeutically.

Prophylactic treatment had no effects under the study execution (**Fig. S5A-F**), however therapeutic treatment showed significant efficacy in multiple measurements. Under the therapeutic protocol, the aged mice were administered the compound orally one day after the SARS-CoV2 infection for 3 of the 5 study days on D1, D2 and D4 and then sacrificed on D5. Therapeutic effects were observed more prominently in the medium dose group of 10 mg/kg, as evidenced by reduced body weight loss over the course of the experiment (**Fig. 5B**), lower lung congestion as assessed macroscopically at the necropsy (**Fig. 5C**), milder lung tissue injury (**Fig. 5D**) and less diffuse alveolar damage (**Fig. 5E**) as evaluated microscopically in the histological examination. The lung tissues from mice in both the 10 and 30 mg/kg groups showed less inflammation and better-preserved alveolar structures (**Fig. 5F**). The viral loads in the lung tissues at the end of the study at D5 were at low levels with no difference among groups (**Fig. S5F**).

The low viral titers at D5 were expected because in this model without treatment, the viral loads in the lung were shown to peak at D2 and to decrease significantly by D4 (36). The viral titer at D5 in this study therefore was less meaningful in terms of antiviral efficacy since the virus in the surviving mice had been largely cleared regardless of treatment. The therapeutic efficacy seen *in vivo*, particularly given that only 3 doses were able to be administered relative to a 5-day course due to site constraints during the pandemic, was very significant both in its endpoints and in its short treatment regimen started after productive viral infection. With its dual actions of host-directed antiviral and antiinflammation, ALD-R491 represents a unique promising oral agent that is worth further investigation as a therapeutic against COVID.

### Efficacy of ALD-R491 in treating and preventing lung injury and fibrosis in a rat model

With the therapeutic efficacy demonstrated in a COVID animal model, we wanted to further evaluate whether the compound of multiple functions could be used to treat or prevent lung injury, and the injury associated lung fibrosis. In COVID-19 patients with pneumonia or requiring intensive care, 42% or over 60% of them developed acute respiratory distress syndrome (ARDS) (37). Interstitial and intra-alveolar fibrosis are hallmarks of ARDS at advanced stages. Up to 25% of ARDS survivors develop physiologic evidence of lung fibrosis within six months. With the continuing rate of SARS-CoV-2 infection worldwide and the increasing populations of long hauler patients, an increase in lung fibrosis is expected in post-COVID patients. Antifibrotic therapies have been proposed for treating severe COVID-19 patients and preventing fibrosis after SARS-CoV-2 infection (38), however the available anti-fibrotic drugs are not well-tolerated and would present especially high risk to COVID-19 patients. Vimentin is the major intermediate filament in fibroblasts, the cell type directly responsible for excessive collagen production and fibrosis. The roles of vimentin in tissue injury and fibrosis have been well-established in the literature. Citrullinated vimentin mediates development and progression of lung fibrosis (39). Vimentin KO or inhibiting vimentin has been shown to decrease collagen production (40), reduce lung injury and protect the lung from developing fibrosis (22, 41).

We tested the vimentin binding compound ALD-R491 in rats with bleomycin-induced lung fibrosis models, using prophylactic as well as therapeutic protocols. In both protocols, ALD-R491 significantly ameliorated lung tissue injury and lung fibrosis. Bleomycin directly injected via intra-trachea into to the unilateral lung caused massive lung injury and severe lung fibrosis. Compared with the positive control treatment with BIBF1120 (Nintedanib), a multiple tyrosine kinase inhibitor marketed as a first line treatment for lung fibrosis, the treatment with ALD-R491 showed significantly improved body weight (**Fig. 6A**), indicating a superior safety profile of ALD-R491. Both therapeutic and prophylactic treatment using ALD-R491 reduced bronchial and arteriole injury in the fibrotic core (**Fig. 6B, 6D**) and the fibrotic border (**Fig. 6E, Fig. S6A),** reduced damage and fibrosis in alveoli (**Fig. S6B, Fig. 6C, 6F**). Interestingly, careful examination revealed that the Masson’s trichrome images for samples with comparable Ashcroft scores exhibited a remarkable difference in connective tissue staining between BIBF1120 and ALD-R491 groups, i.e., ALD-R491 group had a much less intense blue color (stains for fibrosis) than the BIBF1120 group had (**Fig. 6C**). This result indicates that, when assigned comparable Ashcroft scores, the lung tissues from ALD-R491-treated animals had less fibers than the lung tissues from BIBF1120-treated animals, so the Ashcroft scoring on lung fibrosis for ALD-R491 failed to capture this improvement in visible histopathology (less blue) because it falls outside the density rubric of the Ashcroft scoring matrix (scores only density, not color). This discrepancy was further confirmed by collagen-I IHC staining, which showed the actual reduction in fibrosis measured by collagen-I deposition was more significant than that by Ashcroft score in the lung of mice treated by ALD-R491. Compared with vehicle control, both ALD-R491 and BIBF1120 reduced collagen-I deposition in the lung fibrosis core, with the reduction of collagen more significant in ALD-R491 group than in BIBF1120 group (**Fig. 6G, Fig. S6C**). This was consistent with the markable difference in amounts of fibers (blue color) seen by Masson’s staining but not reflected by Ashcroft scores (**Fig. 6C**). Although Ashcroft scoring underestimated the efficacy of ALD-R491 in reducing fibrosis, the results from all the evaluation criteria indicate that ALD-R491 has a superior safety profile and a therapeutic potential not only in ameliorating lung injury but also in preventing and treating lung fibrosis, all of which is directly relevant to use in COVID-19 and its complications.

**Fig. 6.**
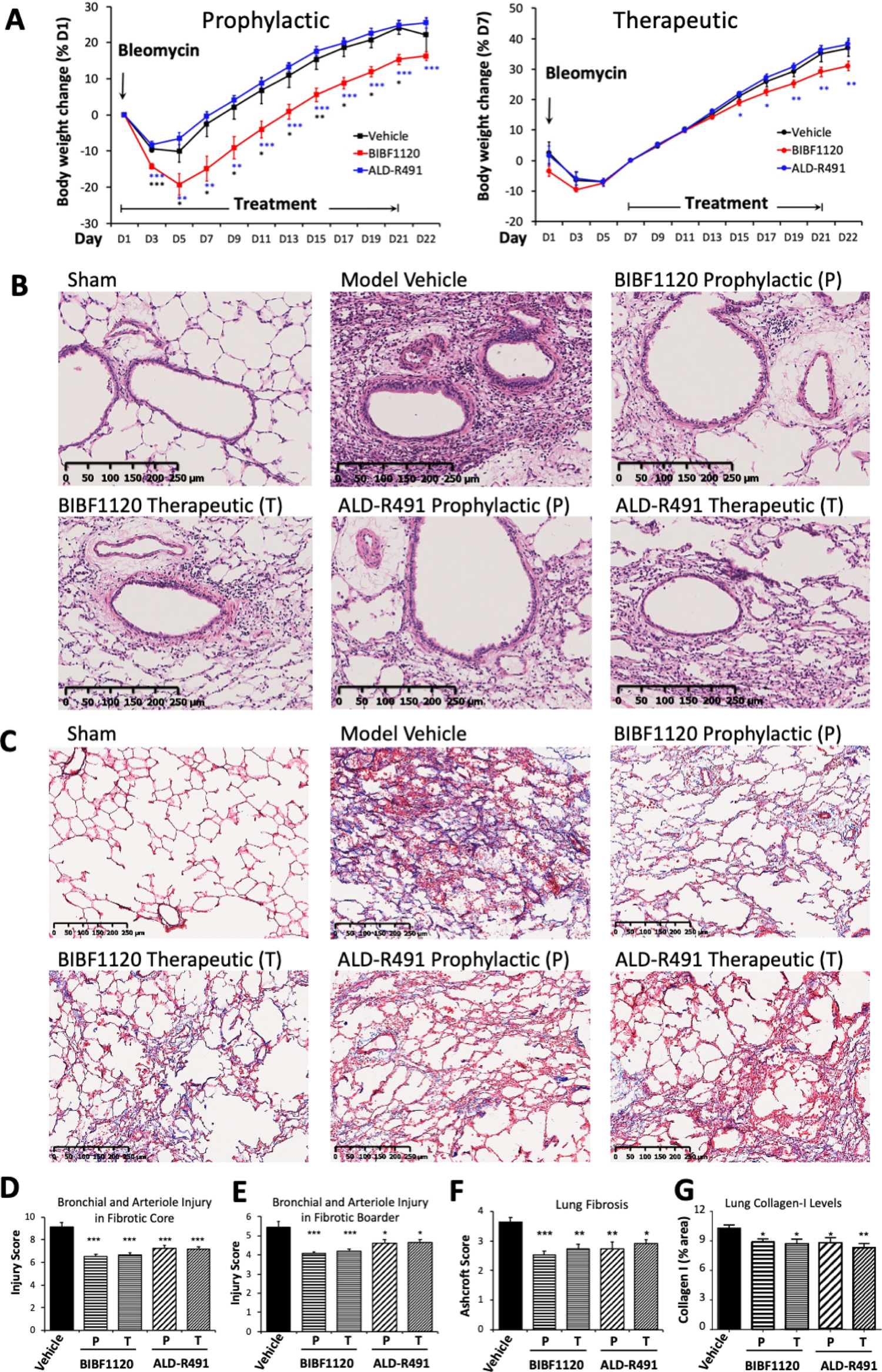
Efficacy of ALD-R491 in treating and preventing lung damage and fibrosis. **A.** Body weight change. Percentage of body weight change was calculated based on the body weight measured on the day when ALD-R491 treatment started. *p <0.05, **p <0.01, ***p<0.001 vs. Model vehicle. **B.** Images for Bronchial and arteriole injury in fibrotic core **C.** Images for alveolar fibrosis. The left lungs were stained with Masson’s trichrome. Blue color: collagen, light red or pink color: cytoplasm, dark brown: cell nuclei. **C.** Bronchial and arteriole injury in fibrotic core. **D-F.** Semi-quantitative scoring of lung injury and fibrosis. Scores for each treatment group were compared with the scores for the model vehicle group. Two-tailed T-test: *p<0.05; **p<0.01; ***p<0.001. n=9. **D.** Bronchial and arteriole injury in fibrotic core. **E.** Bronchial and arteriole injury in fibrotic border. **F.** Lung fibrosis. The Masson’s trichrome stained lung sections were scored according to Ashcraft scoring criteria. **G**. Quantitative measurement of Collagen-1 positive areas in lung fibrotic tissue section. Percentages of Collagen-positive area for each treatment group were compared with that for the model vehicle group. Two-tailed T-test: *p<0.05; **p<0.01; ***p<0.001. n=9.

## DISCUSSION

To address the multifaceted pathogenesis of SARS-CoV2 infection, we are developing a new therapeutic agent with both antiviral and anti-inflammatory action by binding to vimentin (**Fig. 7**). This first-in-class mechanism of action is fundamentally different from that of any existing pipeline agent, or any repurposed drug (13, 42). Importantly, the antiviral mechanisms of ALD-R491 create multiple novel actions against viral infection: first, a blockade of spike protein and ACE2-mediated endocytosis and of endosomal trafficking and exosomal release, which are the cellular processes that are unlikely to be evaded by the virus even with mutations; and second, an enhanced ability of macrophages to kill pathogens, which is critical both for containing the viral spread at early stages and for clearing persistent viral presence after the acute phase of infection. The anti-inflammatory mechanisms of ALD-R491 could also create multiple actions that are novel and different from existing agents, and highly sought for their specific potential against the hyperinflammation of COVID-19: a direct Treg activation, as shown in this study; and a potential inhibition on NLRP3 inflammasome activation, as previous study suggested [22]. The therapeutic effects of the compound on the aged mice with a full-blown, ongoing SARS-CoV2 infections is not only a proxy for its efficacy against severe disease, but also informs its use in the elderly and fragile, two groups of COVID-19 patients with the most concerning unmet need. The therapeutic and preventive effects of the compound on lung injury and fibrosis make it also applicable to patients with post-COVID complications. With oral administration, ALD-R491can be conveniently dosed and broadly accessible to non-hospitalized patients, a population that is growing rapidly as this first pandemic matures.

**Fig. 7.**
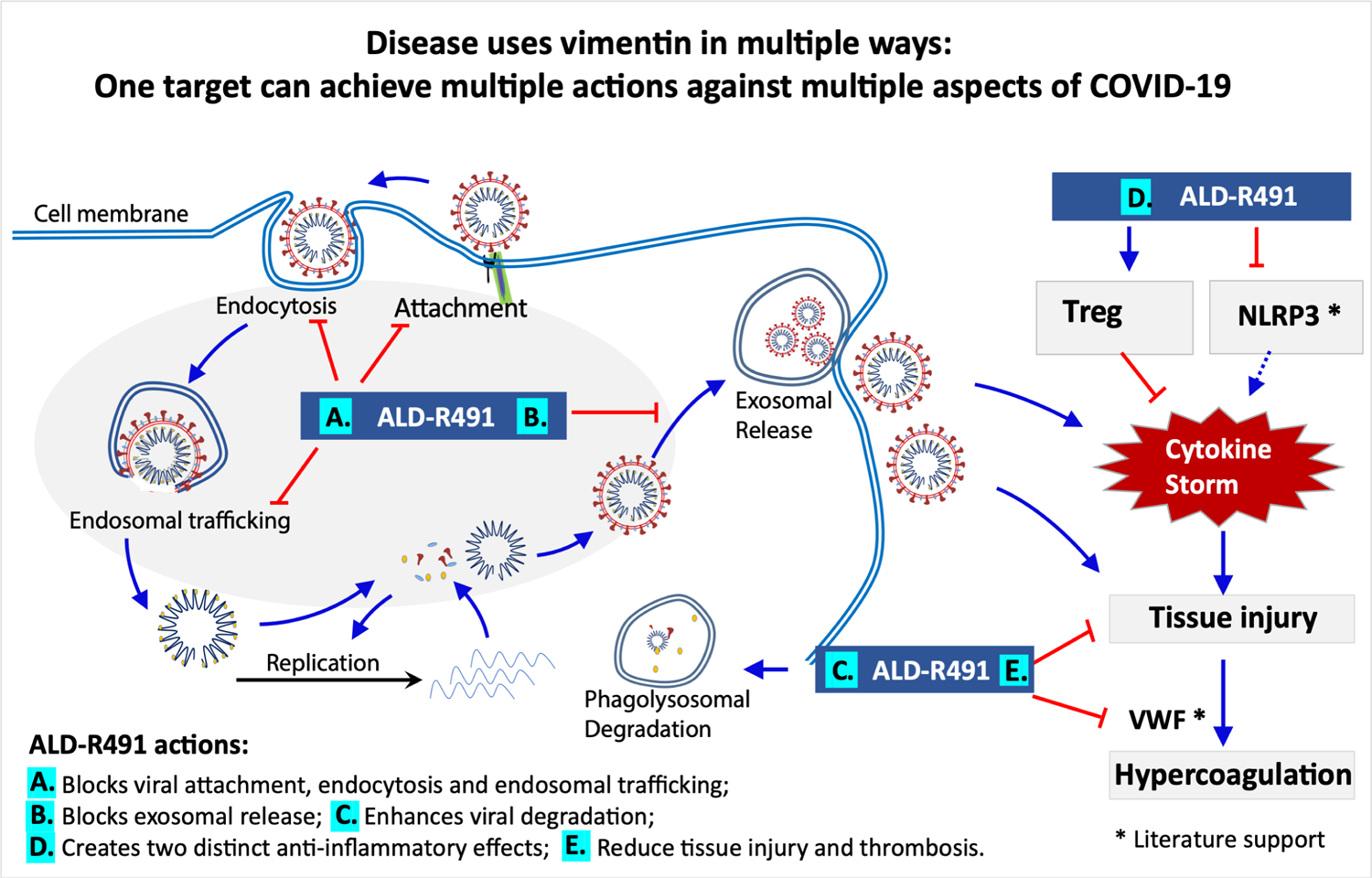
Schematic presentation of actions of ALD-R491 in COVID-19. Left, shows the subcellular events in a cell being infected by a SARS-CoV-2 virus and the host-directed antiviral actions of ALD-R491. **Right,** shows the anti-inflammatory actions of ALD-R491. Both suppressing NLRP3 inflammasome and activating Tregs would dampen cytokine storm and subsequent tissue damage. Curved blue arrows and the shadowed oval area highlight SARS-CoV-2 infection processes. Straight blue arrows indicate positive regulation and red blunt-ended arrows, negative regulation. Treg: Regulatory T cell; NLRP3: NLR family pyrin domain containing 3 inflammasome; VWF: Von Willebrand factor; *: action of vimentin described in literature but not tested in this study.

These remarkable actions can take place because the pathogenesis of SARS-CoV-2 infection involves multiple cellular processes in various types of cells where vimentin is abundantly expressed, including the pulmonary alveolar epithelial cells (AT1 and AT2 cells), vascular endothelial cells, fibroblasts, macrophages and regulatory T cells (43, 44).

The binding of ALD-R491 to vimentin altered the organization and dynamism of the filaments (**Fig. 1A, 1B**), which impacted multiple cellular processes in the cells (**Fig. 1D-F**), leading to a blockade of the initial steps of SARS-CoV-2. Vimentin can function as an attachment receptor on the cell surface and assist membrane and vesicle transport at both anterograde and retrograde directions inside the cell (45, 46). It appears to be widely hijacked by viruses for facilitating their infection, as seen with both SARS-CoV (47) and SARS-CoV2 (48). Our study showed that ALD-R491 potently inhibited Spike protein-ACE2 mediated viral entry through a blockade of endocytosis, the first step of the viral infection (**Fig. 2**). It has been demonstrated that newly assembled SARS-CoV2 particles in the host cell, rather than budding to exit, egress from the cells by fusing multiple vesicle body (MVB)-like structure with the cell membrane, resembling exosomal or lysosomal release pathways (27, 49). Our study showed that vimentin regulated exosomal pathways (25) and that ALD-R491 dose-dependently inhibited exosome release (**Fig. 1F**), which could lead to a blockade of the viral egress. All these actions contribute to a multi-part host-directed antiviral mechanism that offers a multi-part blockage of the transportation machinery hijacked by viruses for productive infection. As the compound acts on cellular processes not the virus itself, it therefore applies to all the SARS viruses regardless of strains or mutations, which is especially advantageous because of the rapid evolution of current SARS-CoV-2 and the potential for future outbreak of other coronaviruses.

ALD-R491’s ability to enhance microcidal activity of macrophages is critical not only for controlling the infection but also for reducing the virus-triggered escalation in inflammation. Monocytes and macrophages are phagocytotic innate immune cells present in the circulation and in the tissues respectively, serving as the first line of defense upon encountering pathogens. Macrophage infiltration in alveoli is a hallmark of COVID-19 (44). Alveolar macrophages express the ACE2 and TMPRSS2 genes that help the cells recognize and internalize SARS-CoV-2. The positive and negative transcripts of SARS-CoV2 have been detected or enriched in alveolar macrophages, monocytes and peripheral blood mononuclear cells (PBMCs) in COVID-19 patients (29, 43), suggesting the existence of live SARS-CoV2 in the monocytes and macrophages. The S1 protein of SARS-CoV-2 has been found persistently present in CD16+ monocytes of post-COVID patients for more than a year (50), suggesting the failure of monocytes to clear the virus. It remains to be investigated whether monocytes and macrophages can support active replication of SARS-CoV2, as has been reported for SARS-CoV and Middle East respiratory syndrome coronavirus (MERS-CoV).

Regardless, monocytes and macrophages may still serve as a permissive system and/or as a viral reservoir, both of which are known to enable virus to spread to other tissues or anchor specifically within the pulmonary parenchyma (51–53), especially when there is a long duration between the onset of symptoms and the development of respiratory failure (6–12 days), or a prolonged course (several months or longer) of syndromes in COVID long-haulers (54). Compromised ability of monocytes and macrophages to clear pathogens in COVID patients might lead to the persistence of the virus inside the cells, escalating a proinflammatory response during the acute phase, and recurring injuries and lingering symptoms in the post-acute phase. Enhancing the ability of macrophages to clear pathogens (**Fig. 3A-B, Fig. S3**), ALD-R491 could therefore be one of ALD-R491’s most distinct, significant, and unique therapeutic benefits in patients with COVID as well as with post-COVID syndromes.

Direct Treg activation a unique feature of ALD-R491 (**Fig. 4, Fig. S4**). The binding of ALD-R491 to vimentin physically disassembles DPC, therefore, directly activates Tregs. This activation occurs rapidly (less than 2 h), unlike other Treg activating mechanisms that require changes in signaling pathway or gene expression, which are indirect and slow acting in nature. For COVID-19 with its rapid disease progression, prompt intervention appears to be especially important, which makes the fast-acting ALD-R491 advantageous. In addition, Treg activation via vimentin targeting was shown not to alter homeostasis of Tregs (55) and therefore would not cause general immunosuppression, which makes it fundamentally different from the use of steroids. Separated from its immune-related functions, Treg activation has been shown to have a major direct and non-redundant role in tissue repair and maintenance (56).

Through directly activating Tregs and other mechanisms, ALD-R491 could not only dampen the overactive immune system without compromising its antiviral response, but also prevent tissues from damage and facilitate their repair. Reduced tissue injury by ALD-R491 has been consistently observed in our studies across different animal models (**Fig. 5-6**).

Anti-fibrosis activity of ALD-R491 is a further additive to a complete view of its impact on COVID-19. Lung pathologies from COVID-19 patients and computed tomography scans of the chest in post-COVID-19 patients showed a significant increase in fibrosis score (44). Fibroblasts are the primary mesenchymal cells in lung tissues.

Overactivation of fibroblasts is the direct cause of lung fibrosis. Alveolar fibroblasts in acute lung injury (ALI) and acute respiratory distress syndrome (ARDS) exhibit a persistent activated phenotype with enhanced migratory and collagen-1 production capacities. Vimentin is the major intermediate filament protein in fibroblasts, promoting the motility and invasiveness of cell and the production of collagen [*40*]. Targeting vimentin has been shown effective in reducing lung fibrosis (22, 41). As lung fibrosis can develop and persist in patients that have recovered from severe COVID-19 (57), the antifibrotic activity of ALD-R491 (**Fig. 6**) should provide additional values against COVID-19 and its complications.

In summary, the vimentin binding compound ALD-R491 impacts multiple cellular processes as well as various types of cells that are involved in the pathogenesis of COVID-19. Its multi-part host-directed antiviral and anti-inflammatory actions and its broad preclinical efficacy and make the compound a promising drug candidate worth further development and exploration for the treatment of COVID-19 and other related diseases.

## MATERIALS AND METHODS

### The compound

The vimentin-binding compound used in this study was ALD-R491, (E) - 1 - (4 - fluorophenyl) - 3 - (4 - (4 - (morpholine - 1 - yl) - 6 - styryl - 1, 3, 5 - triazinyl - 2 - amino) phenyl) urea. It has a molecular formula of C28H26FN7O2 and a purity of > 98%. The compound was synthesized at Bellen Chemistry Co, Ltd., analyzed at Porton Pharma Solutions, Ltd., and provided by Luoda Biosciences, Inc and Aluda Pharmaceuticals, Inc.

### Endocytosis and endosomal trafficking

HEK293T cells were purchased from The Cell Bank of Type Culture Collection of Chinese Academy of Sciences (Shanghai); plasmid PMAX-GFP, from Lonza; transfection agent LipoMAX, from Invitrogen. The cells were grown in 96-well plate to approximately 70% confluence after overnight culture and transfected in 6 replicates with pMAX-GPF plasmid DNA (0.6 μg per well) using LipoMAX according to the manufacture’s instruction. The media were changed at 4 h post transfection, then the culture plates were placed in Incucyte System (Essen Bioscience) for live cell imaging under a 10x objective lens. Fluorescence intensity (GFP signals) in each well was captured once every 2 h continuously till the end of experiment.

### Exosomal release

#### Quantification of exosome release using ExoHTPTM platform

Supernatant (around 98 μl) was collected from each well of the cultured cells, then centrifugated at 300 g for 10 min to remove cells, at 2,000 g for 20 min and 10,000g for 30 min, to remove apoptotic bodies and large extracellular vesicles. Exosomes were isolated from these processed supernatants with ExoEZ^TM^ Exosome Isolation Kit (#ExoCC50, Evomic Science’s, Sunnyvale, CA), according to the manufacturer’s instructions. Luciferase activity in 50 μl of exosome suspension was measured with Promega Renilla Luciferase Assay kit (#E2810, Madison, WI) in TECAN Infinite M100 (TECAN, San Jose, USA).

#### Measurement of nanoparticle size, concentration, and distribution with nanoparticle tracking analysis (NTA)

Overnight cultured Huh7NC12 at 70% confluence were treated for 2 days with media consisting of DMSO and the indicated concentrations of compounds, respectively. Then, culture supernatants were collected, centrifugated at 300 g for 10 min, 2,000g for 20 min, to remove cells and cell debris, apoptotic bodies (Pellet 1), and 10,000g for 30 min, to remove large extracellular vesicles (Pellet 2). Exosomes were isolated from these processed supernatants by ultracentrifugation (100,000 g for 90 min) (Pellet 3). The isolated exosomes from 5 ml or 10 ml of supernatants (Pellet 3) were suspended in 1 ml PBS, visualized on the NanoSight 300 (NanoSight Ltd, Amesbury, UK). The analysis setting was kept constant between samples. The size and number of nanoparticles were calculated.

### Pseudoviral infection

HEK293T/hACE2 cells, Pseudovirus-2019-nCoV-GFP-IRES-LUC and control Pseudovirus-GFP-IRES LUC were purchased from FUBIO (Suzhou). The cells were seeded into 96-well plate at 1×10E4 cells per well, cultured till 40% confluence and mock-infected or infected with the pseudovirus (2×10E7 TFU/mL) at MOI 0.5, 5 and 50, respectively. The cells were exposed to NH4Cl as positive control, DMSO as negative control, and ALD-R491 at concentrations of 0.01, 0.0316, 0.1, 0.316, 1, 3.16, 10 μM for a period from 2 h right before the infection to the start of the infection (Before infection), or for the whole experiment period from 2 h before the infection to the end of the experiment (Full time), or for a period from 2 h after the start of the infection to the end of the experiment (After infection). About 12 h after the infection, the virus-containing media were removed and replaced with each fresh conditional medium, then, the culture plates were placed in Incucyte System (Essen Bioscience) for live cell imaging under a 10x objective lens. Fluorescence intensity (GFP signal) in each well was captured once every 2 h continuously for 48 h, then the cells were measured for luminescence signal by BioTek Synergy 4 plate reader. The viability of mock-infected cells treated with ALD-R491 at different concentrations were measured by MTT assay kit (Beyotime). Each treatment condition was performed in triplicates (n=3).

### Monocyte / macrophage functions

#### The killing of Salmonella by Raw264.7 cells

The cells in a 24-well plate were incubated with ALD-R491 at 1 μM for 2 h. The cells were counted, then added with Salmonella typhimurium (SL1344) to make the ratio of cells to Salmonella 1:10. The culture was gently shaken then centrifuged at 1,000 rpm for 5 min. The cells were incubated in an incubator for 30 min, washed 3 times with PBS, then cultured for 1 h in DMEM medium containing gentamicin at 50 μg/ml. The cells were then lysed with 1% triton X-100, and the lysates were serially diluted and plated on SS plates for 24 h. The bacteria colonies were counted. Six independent experiments were performed.

#### Preparation of Mouse Peritoneal Macrophages

Mice were intraperitoneally injected with 1 ml of 4% Brewer’s modified thioglycolate medium Brewer Modified (BD company, USA). Four days later, the mice were sacrificed with cervical dislocation, soaked in 75% alcohol for 3 min, The abdominal skin was cut open and made the peritoneum fully exposed. The peritoneum was lifted with ophthalmic forceps, injected with 8 ml of pre-cooled PBS into the abdominal cavity. The mouse abdomen was gently massaged for 5 min, and the fluid was withdrawn from the abdominal cavity with a syringe, transferred into a 15 ml centrifuge tube, centrifuged at 4 °C, 1,000 rpm for 10 min. The pelleted cells were resuspended with RPMI 1640 medium and inoculated in a 24-well plate containing round slides. After culturing in an incubator for 2 h, the cells were washed with PBS three times and replaced with a new medium.

#### Dissociation of p47phox and vimentin by ALD-R491 in macrophages

The peritoneal macrophages were incubated in the presence of LPS (5 μg/ml) overnight, and then treated with ALD-R491 (1 μM) for 2 h, fixed with 1% PFA for 10 min, and washed with PBS for three times, permeabilized with 0.2% Triton X-100 for 10 min, washed with PBS for three times, blocked with serum for 30 min and washed with PBS again for three times. The cells were incubated overnight at 4 °C with the anti-p47phox primary antibody (Santa Cruz, sc-17844) and the anti-vimentin primary antibody (Santa Cruz, sc-5565). After washed for three times with PBS, the cells were incubated with the secondary antibody for 1 h in the dark at room temperature. The cells were further stained with DAPI for nuclei, washed with PBS for three times and then mounted. The cells were examined under a confocal microscope and images were taken from three separate fields. Cells with p47phox at locations near cell membrane or/and in the cytoplasm away from cell membrane were counted. The number of the cells with p47phox in the cytoplasm away from the cell periphery was counted from samples treated DMSO or ALD-R491 and percentages of cells with cytoplasmic p47phox were calculated. Three independent experiments were performed.

#### Cellular ROS measurement

Raw264.7 cells were seeded into a 12-well plate and cultured till 90% confluence, then added with ALD-R491 at different concentrations and incubated for another 2 h. DCFH-DA (Beyotime, S0033S) was diluted 1:1,000 with serum-free culture medium to the final concentration of 10 μM, then added to the 12-well plate and incubated for 30 min. The cells were washed twice with PBS, harvested after digestion, and transferred into a 96-well plate. The fluorescence intensity was measured in a microplate reader using the excitation wavelength of 488 nm and the emission wavelength of 525 nm.

### Treg activation

#### Preparation of CD4+CD25+/- cells

CD4+ T cells were isolated from spleen of Balb/c mice (6-8 weeks) by microbeads according to the manufacturer’s instructions. Briefly, single-cell suspension from mouse spleen were incubated with anti-CD4-biotin antibody (Biolegend, 100508) then streptavidin-microBeads (Miltenyi, 130-048-102). Labeled CD4+ cells were positively isolated by LS column and QuadroMACS Separator. Then the CD4+ T cells were stained with anti-CD4-APC (Biolegend, 100412) and anti-CD25-PerCP-CY5.5 (BD, 551071) flow cytometry antibodies. Conventional T cell populations (Tcon, CD4+CD25-) and regulatory T cell populations (Treg, CD4+CD25+) were isolated by BD FACSAria III System.

#### Distal pole complex in Treg cell

The isolated Treg cells (CD4+CD25+) were incubated in RPMI complete culture medium with DMSO or ALD-R491 for 2 h at 37°C. Then the treated Treg cells were collected for Immunofluorescence assay. Briefly, the Treg cells were fixed for 30 min in 4% PFA and permeabilized for 5 min with 0.2% Triton X-100 in PBS. The permeabilized cells were blocked with 3% BSA for 1 h, incubated with anti-Vimentin primary antibody (Proteintech, 10366-1-AP, 1:200) overnight at 4°C. Then, the cells were incubated with AF488 labeled secondary antibody (SouthernBiotech, 4050-30, 1:500), followed by incubation with DAPI (Roche, 10236276001). Each sample was imaged on Nikon A1 confocal microscope for at least three separate fields.

#### Treg function assay

To assess Tregs suppressive function, Tcons were labeled with CFSE (Invitrogen, 65-0850-84) and then co-cultured with Tregs (pretreated with or without ALD-R491) in different ratios in 96-well plate for 3 days. T cells were activated by coated-CD3 antibody (Biolegend, 100302) and diluted-CD28 antibody (Biolegend, 102102) with or without ALD-R491. After 3 days, cells were collected for FACS analysis on BD FACS Canto II. FACS data were analyzed using FlowJo software (TreeStar). The inhibition of Treg cells on Tcon activation was quantified. Three independent experiments were performed.

### Tissue distribution in rats

#### Animals and housing

Sprague-Dawley rats of SPF grades at ages of 6 to 8 weeks were purchased from Beijing Vital River Laboratory Animal Technology Co. Ltd., with animal certificate No. of SCXK (Jing) 2016-0006 and with animal quality certificate No.1100112011030349 and No.1100112011030350. Animals were housed in the animal facility at Joinn Laboratories (Suzhou) with 5 or less rats per gender in each polycarbonate cage and in an environmentally monitored, well-ventilated room (SPF grade) maintained at a temperature of 20 to 26 °C and a relative humidity of 40% to 70%. Fluorescent lighting was provided illumination approximately a 12-hr light/dark cycle per day. Total of 18 rats (9/gender) were used in the study. Only data from 9 female animals (3 animals per time point) are presented.

#### Experiment procedures, data acquisition and analysis

The 9 female rats in 3 groups with body weights from 195 to 214 g were orally administered ALD-R491 at 30 mg/10 ml/kg. At 1, 2, and 6 h after drug administration, one group of the animals were anesthetized and then euthanized using abdominal aorta exsanguinations, and their blood and tissue samples were taken. The concentrations of ALD-R491 in rat plasma, tissue, feces, urine, and bile samples were analyzed by the validated LC-MS/MS methods with the LLOQ (Lower Limit of Quantification) concentrations of 0.500 ng/mL for plasma, 0.500 ng/mL (equal to 25.0 ng/g after conversion) for tissue. The concentration value below LLOQ was set as 0 for the purpose of calculation. Software of LabSolutions (Version 6.81 SP1) from SHIMADZU Company was used for outputting raw chromatograms and generating concentration and accuracy results of the plasma samples. WinNonlin was used for distribution data statistical analysis (including mean, SD, and %CV) of plasma and tissues concentrations. Microsoft Office Excel (2007) was used for tissue concentration column graphs.

### SARS-CoV-2 M10 infection in aged mice

#### Animals and housing

Aged (11- to 12-month-old) female BALB/c mice were obtained from Envigo (retired breeders). The animals were housed in the animal facility at University of North Carolina, Chapel Hill, with 5 mice in each cage with food and water provided ad libitum, with a 12 h/12 h light/dark cycle. The animals were acclimated for 7 days in the BSL-3 facility prior to any experimentation

#### Virus and model establishment

Baric laboratory generated the stock of SARS-CoV-2 MA10, a mouse-adapted virulent mutant created from a recombinantly derived synthesized sequence of the Washington strain. The virus was maintained at low passage (P2-P3) to prevent the accumulation of additional potentially confounding mutations. Mice were anesthetized i.p. with a combination of 50 mg/kg Ketamine and 15 mg/kg Xylazine in 50 μl, then infected i.n. with 10^3^ PFU of sequence- and titer-verified SARS-CoV-2 MA10 in 50 μl of DMEM diluted in PBS to the inoculation dosage

#### Experiment groups and treatment regimens

Eighty animals were used in this study., Ten animals per group were randomly assigned in 1 mock-infected vehicle treatment group and 7 virus (10^3^ PFU SARS-CoV-2 MA10)-infected groups treated p.o. via oral gavage respectively with vehicle and ALD-R491 at 3, 10 and 30 mg/kg in prophylactic protocol (dosing at −15, −5, +24, +48, +72, +96 h post infection) or therapeutic protocol (dosing at +24, +48, +96 h post infection).

#### Procedure and endpoint

Daily clinical evaluation and scoring were performed including body weight and symptom. The animals that survived anesthesia were carried to the experimental endpoint. As the mice were continuing to lose weight at all doses and in both treatment arms and the disease scores were increasing, all animals were euthanized at 5 days post-infection. Euthanasia was performed by inhalational isoflurane (drop method) and thoracotomy, with removal of vital organs (lungs). Lung tissue was taken for assessments of titer, RNA, and histology. Serum was taken for downstream analysis.

#### Histopathological examination of the lung

Lung specimens were fixed in formalin and stained with hematoxylin and eosin for histological assessments. Lung samples were scored by a board-certified veterinary pathologist for acute lung injury (ALI) and diffuse alveolar damage (DAD). The ATS ALI score is a weighted composite score based on the Matute-Bello rubric and considers neutrophils in the alveolar space, neutrophils in the interstitium, hyaline membrane formation, protein in airspaces, and alveolar septal thickening. All lung specimens were submitted for scoring; the pathologist scored all specimens that were deemed suitable. Thus, at least 6 lungs were scored in all conditions.

**Statistical analyses** were performed by Kruskal-Wallis test (Titer), One-way ANOVA (Congestion score) or two tailed T Test (Weight loss, ATS ALI and DAD scores).

### Bleomycin-induced lung injury and fibrosis in rats

#### Animals and housing

Forty-five male SD rats of SPF grade were purchased from Beijing Vital River Laboratory Animal Technology Co., Ltd. with the company certificate No.: SCXX (Jing) 2012-001 and the animal use certificate No. 33000800003683. The animals were housed in the Animal House Facility of the KCI Biotech (Suzhou) Inc. at a temperature of 20 to 26°C and a relative humidity of 40% to 70%. Fluorescent lighting was provided illumination approximately a 12 h light/dark cycle per day. The experimental protocol was reviewed and approved by the Institutional Animal Care and Use Committee (IACUC) at KCI Biotech Inc. All experimental procedures were conducted in conformity with the institutional guidelines issued by KCI Biotech Inc.

#### Model Establishment

This study was carried out in strict accordance with the SOP institutional guidelines for the care and use of laboratory animals. The rats were anesthetized by intraperitoneal injection of pentobarbital sodium at dose of 50mg/kg. The skin in the neck area was disinfected, then the skin layers were carefully incised to have the trachea exposed. Bleomycin (BLM) was directly injected into left main bronchus at a dose of 3mg/kg body weight in volume of 1.0 ml/kg via a cannula. After the skin layers were closed, the animals were placed on an electric heat pad at 37 °C until they woke up from anesthesia before returning to holding cages with free access to water and diet.

#### Experiment Groups and treatment regimen

The modeled rats were randomly assigned into 5 groups (n=9 per group), treated p.o. via oral gavage respectively with vehicle at 5 ml/kg/day, with BIBF1120 at 100 mg/kg/d prophylactically (q.d., from Day 1 to Day 21) or therapeutically (q.d., from Day 8 to Day 21), and with ALD-R491 at 100 mg/kg/d prophylactically (q.d., from Day 1 to Day 21) or therapeutically (q.d., from Day 8 to Day 21). One day after the last dosing (Day 22), all animals were euthanized according to the standard SOP at KCI Biotech. The animals were perfused systemically with ice-cold saline, their left lungs were harvested and perfuse-fixed with 10% formaldehyde solution (3ml of each lung), then processed for pathological analysis.

#### Histopathological examination of the left lungs

The whole left lung was dehydrated and embedded in paraffin wax. The sections were cut at a thickness of 4um and processed for H&E and Masson’s Trichrome staining. All the procedures were performed according to KCI pathology SOPs. Whole slides were then scanned by Hamamatsu NanoZoomer Digital Pathology S210 slide scanner after staining.

Bronchiole and pulmonary arteriole damage and inflammatory cell infiltration in fibrosis core and fibrosis boarder areas were scored with H&E-stained slides.

Criteria for grading the damage of terminal bronchiole wall were: score 0 for normal structure; score 1 for normal structure with less than 1/2 of the terminal bronchiole wall area injury and characterized by bronchial epithelial cells damage and epithelium regeneration, wall edema, medium layer of the mucosal muscle degeneration or regeneration; score 2 for structure with more than 1/2 of the terminal bronchiole wall area injured and characterized by bronchial epithelial cells damage and epithelium regeneration, wall edema, medium layer of the mucosal muscle degeneration or regeneration; score 3 for structure with more than 1/2 area of the terminal bronchiole wall injured and characterized by bronchial epithelial cells damage and epithelium regeneration, wall edema, medium layer of the mucosal muscle degeneration or regeneration, granulomas formation or fibrosis.

Criteria for grading inflammatory cell infiltration in the terminal bronchiole wall were: score 0 for normal structure with no inflammatory cells infiltration; score 1 for a few scattered but localized inflammatory cell infiltration (less than 10 cells per field) in the outer wall of the terminal bronchiole; score 2 for numerous scattered inflammatory cell infiltration that was focal or diffuse, affecting less than 1/2 of the total area of the terminal bronchiole wall; .score 3 for diffuse infiltration of inflammatory cells that affected more than 1/2 of the total area of the terminal bronchiole wall and extended to the inner and the medium layers of basement membrane.

Criteria for grading the damage of pulmonary small arteries wall were: score 0 for the clear and complete structure of pulmonary small arteries; score 1 for partially exfoliated endothelial cells; score 2 for exfoliated endothelial cells with degeneration, regeneration or small focal necrosis in the medium layer of smooth muscle; score 3 for exfoliated endothelial cells with degeneration, regeneration or small focal necrosis and formation of granulomas or fibrosis.

Criteria for grading inflammatory cell infiltration in the pulmonary arteriole were: score 0 normal structure of pulmonary small arteries; score 1 for a few scattered and localized inflammatory cell infiltration (less than 10 cells per field) in the outer wall of pulmonary small arteries; score 2 for numerous scattered inflammatory cell infiltration, affecting less than 1/2 of the total area of outer wall of pulmonary small arteries; score 3 for diffuse infiltration of inflammatory cells that affected more than 1/2 of the total area of 1/2 area of the pulmonary small artery wall and extended to the medium layer of the basement membrane.

BLM-induced left lung injury and fibrosis were evaluated with Masson Trichrome stained slides using Ashcroft scoring matrix. The criteria were: score 0 for the normal structure with no fibrotic lesion in alveolar septum; score 1 for the isolated and simple pulmonary fibrosis in alveolar septum (alveolar walls thicken but thinner than triple of that for a normal lung), and little exudate and no fibrosis material in major alveolar areas; score 2 for the clear fibrosis change in alveolar septum (alveolar walls thicker than triple of that for a normal lung) with formation of small nodules that were not connected, and little exudate and no fibrosis material in major alveolar areas; score 3 for the formation early stage fibrosis forms in all alveoli in alveolar septum (alveolar walls thicker than triple of that for a normal lung), and little exudate and no fibrosis material in major alveolar areas; score 4 for the lung fibrosis with alveolar septum still visible, and the formation of isolated fibrosis nodules in alveolar areas (≤10% at high magnification); score 5 for the lung fibrosis with alveolar septum still visible, the formation of integrated fibrosis nodule in alveolar areas (>10% and ≤50% at high magnification), and the lung structure substantially impaired but still existed; score 6 for the seen but barely alveolar septum existed, and the formation of integrated fibrosis nodule in alveolar areas (>50% at high magnification), and the lung structure barely existed; score 7 for the disappeared alveolar septum, and the pulmonary alveoli and interstitial fibrosis proliferation seen but still with vacuole structures; score 8 for the disappeared alveolar septum, and the pulmonary alveoli and interstitial fibrosis proliferation seen at high magnification. **immunohistochemistry (IHC) for Collagen-I:** The IHC staining was processed according to the standard IHC protocol at KCI Biotech. Anti-collagen-I antibody (Abcam, Cat# ab34710) was used. The stained slides were then scanned by Hamamatsu NanoZoomer Digital Pathology S210 slide scanner and analyzed using the software to calculate the positive staining area/analysis area (%) or positive staining cells per unit area.

## Statistical analyses

Statistical analysis was performed using Graphpad prism 6.0 software. Descriptive results were expressed as mean ± sem or mean ± sd. Statistical comparisons were performed using t-test, one-way analysis of variance or two-way analysis of variance test. *p<0.05* was considered as statistically significant.

## Author contributions

RC conceived the studies; YX and RC designed the studies; YX and RC supervised and ZL, JW, JZ, LM, BY, ZQ, FZ, YD, KH, ZWL and TW performed the in vitro studies. ZL, JW, JZ and BY contributed equally to this work. ZL, BY, YD performed transfection and pseudoviral infection experiments. JW, LM performed exosomes experiments and pilot Treg experiments. ZL, KH performed macrophage experiments. ZL and KH performed Salmonella killing assays in R. Huang’s lab at Soochow University. FZ performed immunofluorescence experiments. ZQ performed the FRAP experiments. ZWL performed cell preparation from animals for in vitro studies. JPZ supervised and JZ performed Treg cell experiments. ES supervised and JC performed the lung fibrosis *in vivo* studies, JG supervised and WW and performed *in vivo* drug tissue distribution studies, DS managed the SARS-CoV2 *in vivo* project that was performed in R. Baric’s lab at University of North Carolina. RC wrote the manuscript. All authors read, edited and approved the final version of the manuscript.

## Materials & Correspondence

Correspondence and material requests should be addressed to Ruihuan Chen at, or Yong Xu at. All data are available in the main text or the supplementary materials. The compound ALD-R491 is available under a materials transfer agreement.

## SUPPLEMENTARY FIGURE LEGENDS

**Fig. S1. ALD-R491 has no effect on other type III intermediate filaments.** Left. No structural changes in GFAP in human glioma U87 cells treated with or without the compound ALD-R491. Right. No structural changes in desmin in human osteosarcoma U2OS cells treated with or without the compound ALD-R491.

**Fig. S2. ALD-R491 blocked Spike protein-ACE2 mediated viral entry in vitro.** Upper. Time course of the GFP reporter gene expression in the ACE2-overexpressing HEK293 cells infected with the SARS-CoV spike protein-pseudotyped-lentivirus at MOI of 5. **Lower.** Dose-dependent reduction of the GFP expression at the time point of 48 hrs post viral infection.

**Fig. S3. Schematic presentation of effects of ALD-R491 on macrophage.** ALD-R491. The binding of ALD-R491 to vimentin releases p47phox, leading to the activation of NADPH oxidase in the phagosome, and therefore an increase of pathogen-killing capacity of macrophage.

**Fig. S4. Schematic presentation of effects of ALD-R491 on Treg cell.** The binding of ALD-R491 to vimentin disrupts the distal pole complex of Treg cell, consequently, allow the suppressive molecules move to the immunological synapse, through which Treg cell is directly activated.

**Fig. S5. Prophylactic treatment with ALD-R491 had no effect on the aged mice with SARS-CoV2 M10 infection.** A. Body weights were measured daily over the course of study. Body weight changes were calculated based on the body weight prior to the infection. **B.** Lung congestion was evaluated on Day 5. **C.** ATS Acute Lung Injuries (ALI) were scored by the method of Matute-Bello (2011). **D.** The diffuse alveolar damages (DAD) were scored by the method of Schmidt (2018). **E**. Histopathology was blindly evaluated by a board-certified pathologist. **F**. Lung tissues from all groups of mice were harvested on Day 5 and viral titers were measured by PCR. No statistical significance between infected vehicle group vs. individual treatment groups.

**Fig. S6. ALD-R491 reduced lung injury and collagen deposition in the lung.** Representative images of **A.** the bronchial and arteriole injury in fibrotic border; **B.** the alveolar damage in fibrotic core; **F.** the immunohistochemical staining of collagen 1 in the fibrotic lung. Brown color: collagen 1. All the representative images were selected from sections with scores or percentage of positive areas close to those of individual group averages.

